# Omnipose: a high-precision morphology-independent solution for bacterial cell segmentation

**DOI:** 10.1101/2021.11.03.467199

**Authors:** Kevin J. Cutler, Carsen Stringer, Paul A. Wiggins, Joseph D. Mougous

## Abstract

Advances in microscopy hold great promise for allowing quantitative and precise readouts of morphological and molecular phenomena at the single cell level in bacteria. However, the potential of this approach is ultimately limited by the availability of methods to perform unbiased cell segmentation, defined as the ability to faithfully identify cells independent of their morphology or optical characteristics. In this study, we present a new algorithm, Omnipose, which accurately segments samples that present significant challenges to current algorithms, including mixed bacterial cultures, antibiotic-treated cells, and cells of extended or branched morphology. We show that Omnipose achieves generality and performance beyond leading algorithms and its predecessor, Cellpose, by virtue of unique neural network outputs such as the gradient of the distance field. Finally, we demonstrate the utility of Omnipose in the characterization of extreme morphological phenotypes that arise during interbacterial antagonism and on the segmentation of non-bacterial objects. Our results distinguish Omnipose as a uniquely powerful tool for answering diverse questions in bacterial cell biology.

## Introduction

Although light microscopy is a valuable tool for characterizing cellular and sub-cellular structures and dynamics, quantitative analysis of microscopic data remains a persistent challenge^1^. This is especially pertinent to the study of bacteria, many of which have dimensions in the range of optical wavelengths. Thus, their cell body is composed of a small number of pixels (*e.g*., ~100-300 px^2^ for *E. coli* at 100x magnification). At this scale, accurate subcellular localization requires defining the cell boundary with single-pixel precision. The process of defining cell boundaries within micrographs is termed cell segmentation and this is a critical first step in current image analysis pipelines^2,3^.

In addition to their small size, bacteria adopt a wide range of morphologies. Although many commonly studied bacteria are well-approximated by idealized rods or spheres, there is growing interest in bacteria with more elaborate shapes^4^. Some examples include Streptomycetales, which form long filamentous and branched hyphal structures^5^, and Caulobacterales, which generate extended appendages distinct from their cytoplasm^6^. Furthermore, microfluidic devices are allowing researchers to capture the responses of bacteria to assorted treatments such as antibiotics, which often result in highly irregular morphologies^7^. Whether native or induced, atypical cell morphologies present a distinct problem at the cell segmentation phase of image analysis^8,9^. This is compounded when such cells are present with those adopting other morphologies, as is the case in many natural samples of interest^10^. To date, there are no solutions for segmenting bacterial cells of assorted shape and size in a generalizable manner^1^.

Cell segmentation is a complex problem that extends beyond microbiological research, thus many solutions are currently available in image analysis programs^8,9,11–27^. Most of these solutions use traditional image processing techniques such as the application of an intensity threshold to segment isolated cells; however, this approach does not perform well for cells in close contact and it requires extensive parameter-tuning in order to optimize for a given cell type. SuperSegger was developed to address these issues specifically in bacterial phase contrast images^13^. This program utilizes both traditional image filtering techniques and a shallow neural network to correct for errors that thresholding and watershed segmentation tend to produce.

Deep neural networks (DNNs) are now widely recognized as superior tools for cell segmentation^28^. Unlike traditional image processing, machine-learning approaches such as DNNs require training on a ground-truth dataset of cells and corresponding labels. Trained DNNs are thus limited in applicability to images that are representative of those in the training dataset. Early DNN approaches were based on the Mask R-CNN architecture^24^, whereas more recent algorithms such as StarDist, Cellpose, and MiSiC are based on the U-Net architecture^12,15,26^. Pachitariu and colleagues showed that Cellpose outperforms Mask R-CNN and StarDist on a variety of cell types and cell-like objects, distinguishing it as a general solution for cell segmentation^12^. Notably, the representation of bacteria in their study was limited. MiSiC was developed as a general DNN-based solution for bacterial segmentation; however, the authors of MiSiC did not provide comparisons to other DNN algorithms^15^. Here, we evaluated the performance of state-of-the-art cell segmentation algorithms on a diverse collection of bacterial cells. Our findings motivated the design of a new algorithm, Omnipose, that significantly outperforms all previous cell segmentation algorithms across a wide range of bacterial cell sizes, morphologies, and optical characteristics. We have made Omnipose and all associated data immediately available to researchers, and we anticipate that our model – without retraining – can be applied to diverse bacteriological systems. Furthermore, following the incorporation of additional ground truth data, Omnipose could serve as a platform for segmenting various eukaryotic cells and extended, anisotropic objects more broadly.

## Results

### Evaluation of bacterial cell segmentation algorithms

Numerous image segmentation algorithms have been developed, and the performance of many of these on bacterial cells is documented^1^. These broadly fall into three categories: (*i*) traditional image processing approaches (*e.g*., thresholding, watershed), (*ii*) traditional/machine learning hybrid approaches, and (*iii*) deep neural network (DNN) approaches. Given the goal of developing software with the capacity to recognize bacteria universally, we sought to identify strongly performing algorithms for further development. An unbiased, quantitative comparison of cell segmentation algorithms on bacterial cells has not been performed; thus, we selected one or more representatives from each category for our analysis: Morphometrics^23^ (*i*), SuperSegger^13^ (*ii*), Mask R-CNN^27^, StarDist^26^, MiSiC^15^, and Cellpose^12^ (*iii*). For a detailed review of these choices, see Methods.

For training and benchmarking these algorithms, we acquired micrographs of assorted bacterial species representing diverse morphologies and optical characteristics. Many studies of bacteria involve mutations or treatments that cause extreme morphologies. To capture this additional diversity, we included wild-type and mutant bacteria grown in the presence of two beta-lactam antibiotics, cephalexin and aztreonam, and A22, which targets MreB^29^. Finally, based on our interest in microbial communities, we acquired images of mixtures of bacteria which display distinct morphologies and optical characteristics. In total, we collected 4833 images constituting approximately 700,900 individual cells deriving from 14 species (Extended Data Table 1). Next, we developed a streamlined approach for manual cell annotation and applied it to these images (see Methods), yielding 46,000 representative annotated cells that serve as our ground-truth dataset. We arbitrarily split this data into a 27,000-cell training set and a 19,000-cell benchmarking set. Relevant cellular metrics did not differ substantially between the groups, confirming that the benchmarking set faithfully represents the training set (Extended Data Fig. 1).

To facilitate direct comparison of the algorithms, we first optimized their performance against our data. For the DNN approaches, each algorithm was trained on our dataset using developer-recommended parameters. Morphometrics and SuperSegger cannot be automatically optimized using ground-truth data; therefore, we manually identified settings that optimized the performance of these algorithms against our dataset (see Methods). As a quantitative measure for algorithm performance, we compared their average Jaccard Index (JI) as a function of intersection over union (IoU) threshold – a well-documented approach for evaluating image segmentation (Fig. 1a)^30,31^. IoU values lie between zero and one, with values greater than 0.8 marking the point at which masks become indistinguishable from ground truth by the expert human eye (Extended Data Fig. 2)^30^. This analysis showed that DNN-based approaches significantly outperform other algorithms. However, within the DNN group, substantial differences in performance were observed; Cellpose and StarDist outperform Mask R-CNN and MiSiC at high IoU thresholds. The performance of all algorithms varied greatly across the images in our ground-truth dataset, with much of this variability delineated by cell type and morphology categories (Fig. 1b-g). Whereas all other algorithms exhibited visible segmentation errors in two of the three cell categories we defined, errors by Cellpose – the best overall performing algorithm at high IoU thresholds – were only apparent in elongated cells (Fig. 1h-j).

**Fig. 1.**
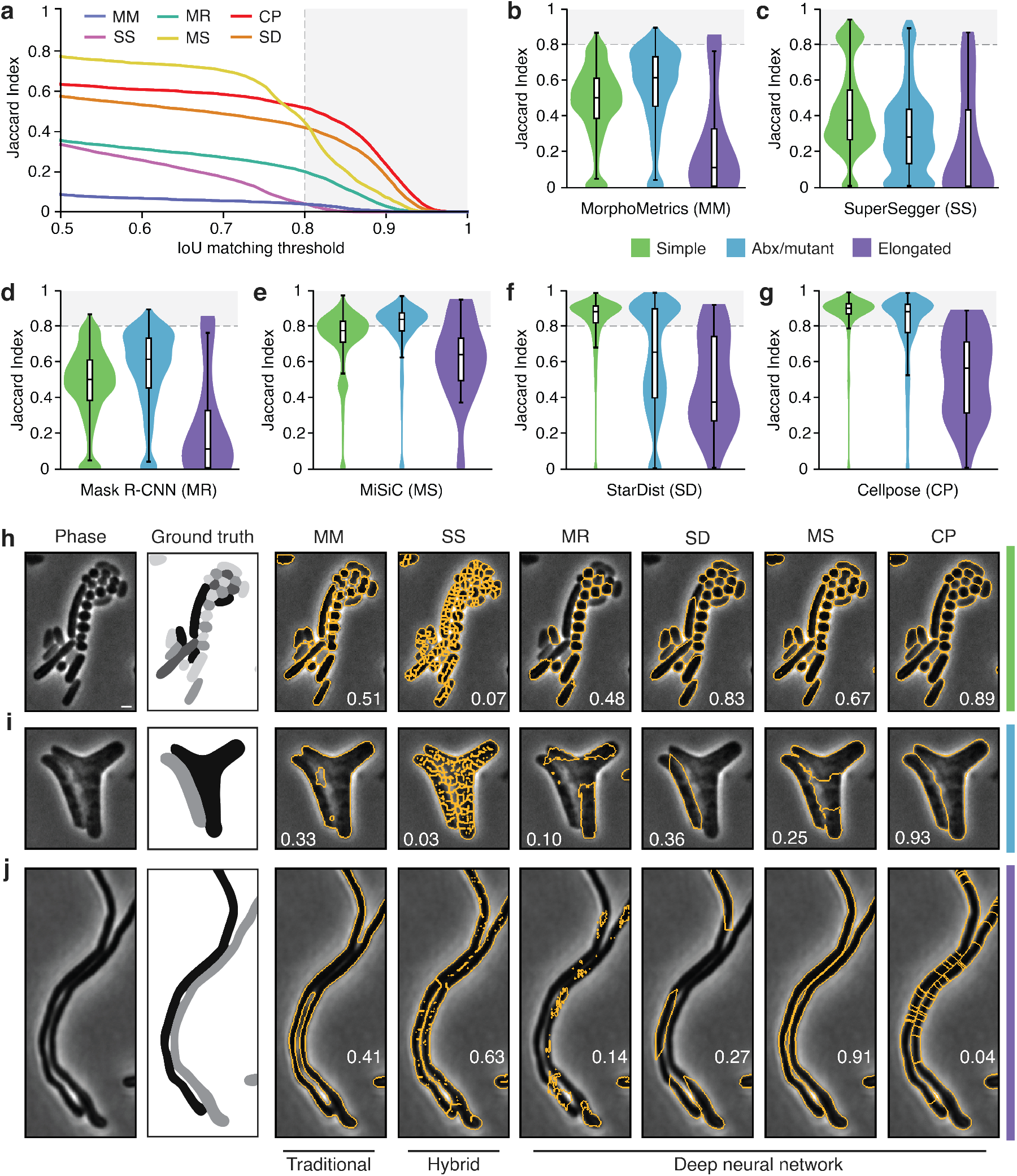
Quantitative comparison of segmentation methods distinguishes Cellpose as a high performing algorithm. (**a-g**) Comparison of segmentation algorithm performance on our test dataset. (**a**) Overall performance measured by Jaccard Index (JI). The JI was calculated at the image level and values averaged across the dataset are displayed. Algorithm abbreviations defined in B-G. (**b-g**) Algorithm performance partitioned by cell type (Simple, n=12,869; Abx/mutant, n=6,138; Elongated, n=46). Images were sorted into types as defined in Supplemental Table 1 (Abx, antibiotic). (**h-j**) Representative micrographs of cell type partitions analyzed in B-G, indicated by vertical bars at right. Ground-truth masks and predicted mask outlines generated by the indicated algorithm are displayed. Mean matched IoU values for cells shown are displayed within each micrograph. Bacteria displayed are (H) *Vibrio cholerae, Pseudomonas aeruginosa, Bacillus subtilis, Staphylococcus aureus*, (I) aztreonam-treated *Escherichia coli* CS703-1, and (J) *Streptomyces pristinaespiralis*. All images scaled equivalently; scale bar is 1mm.

### Motivation for a new DNN-based segmentation algorithm

Our comparison revealed that Cellpose offers superior performance relative to the other segmentation algorithms we analyzed, and for this reason, we selected this algorithm for further development. Notably, even at the high performance levels of Cellpose, only 83% of predictions on our benchmarking dataset are above 0.8 IoU. This limits the feasibility of highly quantitative studies such as those involving subcellular protein localization or cell–cell interactions.

Cellpose utilizes a neural network that is trained on ground-truth examples to transform an input image into several intermediate outputs, including a scalar probability field for identifying cell pixels (Extended Data Fig. 3a, panels *i-iii*)^12^. Cellpose is unique among DNN algorithms by the addition of a vector field output, which is defined by the normalized gradient of a heat distribution from the median cell pixel coordinate (Extended Data Fig. 3a, panels *iv,v*). This vector field directs pixels toward a global cell center via Euler integration, allowing cells to be segmented based on the points at which pixels coalesce (Extended Data Fig. 3b). In contrast to other algorithms, this approach for reconstructing cells is size- and morphology-independent, insofar as the cell center can be correctly defined.

To further interrogate the accuracy of Cellpose on our dataset, we evaluated its performance as a function of cell size. We compared cell area against the number of segmentation errors, calculated as the number of redundant or missing masks corresponding to each ground-truth cell mask. This revealed a strong correlation between cell size and segmentation errors, with the top quartile of cells accounting for 83% of all errors (Fig. 2a). To understand the source of these errors, we inspected the flow field output of many poorly segmented cells across a variety of species and growth conditions. This showed that elongated cells, an important morphology often seen in both wild-type and mutant bacterial populations, are particularly susceptible to over-segmentation (Fig. 2b). We attribute this to the multiple sinks apparent in the corresponding flow fields. In the Cellpose mask reconstruction algorithm, pixels belonging to these cells are guided into multiple centers per cell, fragmenting the cell into many separate masks.

**Fig. 2.**
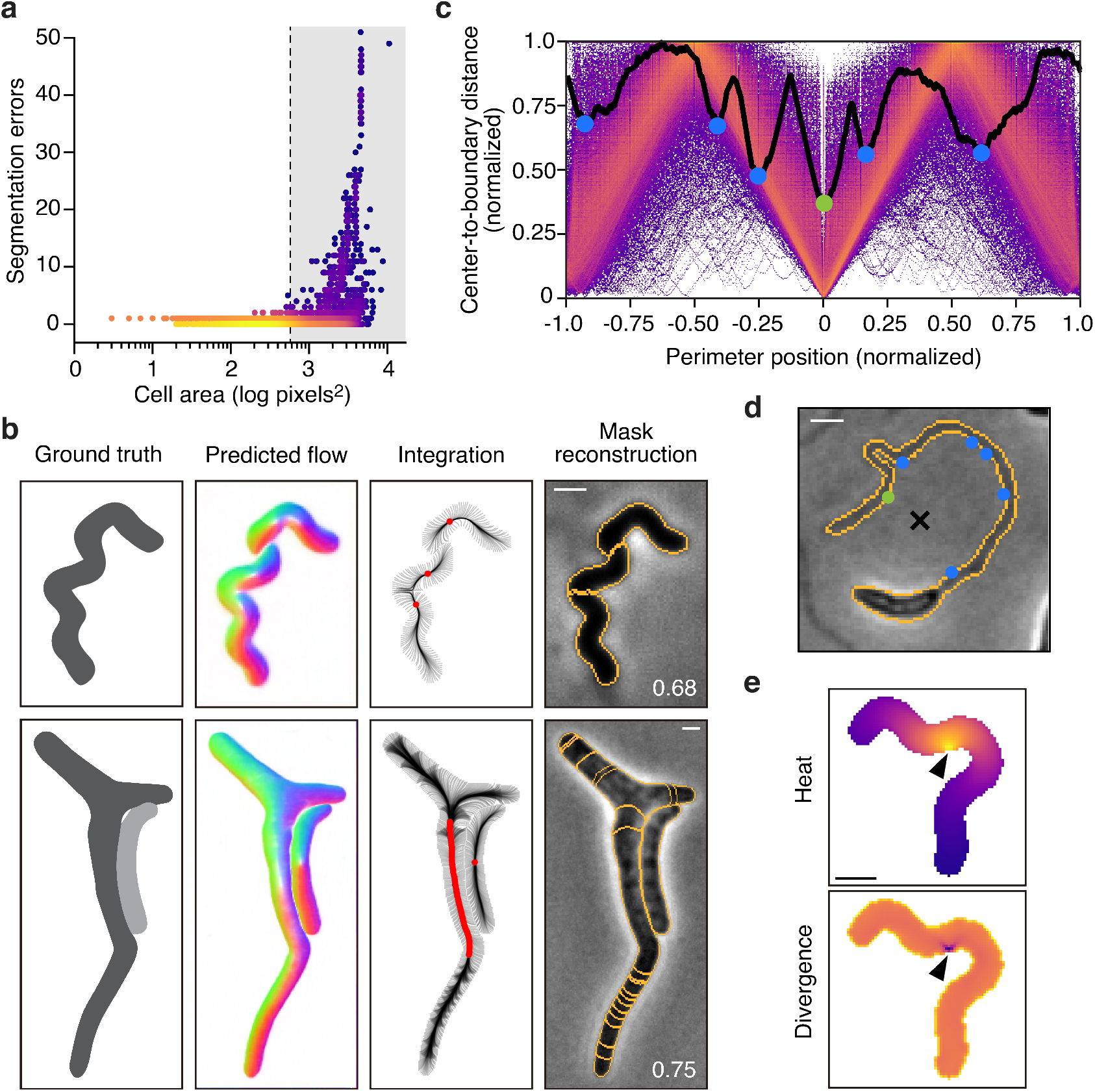
Cellpose over-segments extended, anisotropic cells. (**a**) Single-cell analysis of segmentation error as a function of cell area. Color represents density on a log scale. Gray box represents the top quartile of cell areas. (**b**) Representative examples exhibiting problematic flow fields. Corresponding boundary pixel trajectories are shown in black and final pixel locations in red. Predicted mask overlays are shown with mean matched IoU values. (**c**) Analysis of stochastic center-to-boundary distances. Distance from the center (median pixel coordinate) to each boundary pixel is normalized to a maximum of 1. Position along the boundary is normalized from −1 to 1 and centered on the point closest to the median pixel. Center-to-boundary for the cell in panel D is highlighted in black. (**d**) Representative cell with median coordinate outside the cell body (black X). Cellpose projects this point to the global minima of this function (green dot), but several other local minima exist (blue dots). (**e**) The heat distribution resulting from a projected cell center (black arrow). The normalized gradient corresponds to the divergence shown. Bacteria displayed are (a,e) *Helicobacter pylori*, (b) *Escherichia coli* CS703-1, both treated with aztreonam, and (d) *Caulobacter crescentus* grown in HIGG media. Scale bars are 1 μm.

We hypothesized that the observed defect in Cellpose flow field output is a consequence of two distinct flow field types arising from our training dataset: those where the median pixel coordinate, or ‘center’, lies within the cell (97.8%) and those where it lies outside the cell (2.2%). In the latter, Cellpose projects the center point to the nearest boundary pixel, ultimately leading to points of negative divergence on the cell periphery that are chaotically distributed (Fig. 2c-e). On the contrary, non-projected centers maintain a uniform field magnitude along the entire boundary and adhere to the global symmetries of the cell (Extended Data Fig. 4a,d). A similar issue is also encountered in cells with centers that are not projected but lie close to the boundary (Extended Data Fig. 4b-d). Cells with a center point closer than 0.3 times the mean cell diameter (a factor of 0.2 off-center) to the boundary account for an additional 8.5% of our data. Neural networks can be exquisitely sensitive to the outliers in their training data^32^; therefore, we suspect that this small fraction of corrupt flow fields has significantly impacted the performance of Cellpose.

### Development of a new DNN-based segmentation algorithm

As there exists no straightforward means of defining a cell center for irregular objects, we sought to develop a segmentation algorithm that operates independently of cell center identification. We built our new algorithm, which we named Omnipose, around the scalar potential known as the distance field (or distance transform), which describes the distance at any point 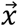 in a bounded region Ω to the closest point on the boundary *∂*Ω. Notably, this widely utilized construct is one of the intermediate outputs of StarDist^32^. Whereas in StarDist it is used to seed and assemble star-convex polygons, its use in Omnipose is to define a new flow field within the Cellpose framework. The use of a distance field has several advantages. First, the distance field is defined by the eikonal equation 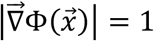, and so its gradient has unit magnitude throughout the bounded region for which it is calculated. This grants it faster convergence and better numerical stability when compared to alternative solutions producing similar fields (*e.g*., screened Poisson; see Methods) (Extended Data Fig. 5a). Second, the distance field is independent of morphology and topology, meaning that it is applicable to all cells. Lastly, the resulting flow field points uniformly from cell boundaries toward the local cell center, coinciding with the medial axis, or skeleton, that is defined by the stationary points of the distance field (Extended Data Fig. 5b). This critical feature allows pixels to remain spatially clustered after Euler integration, solving the problem of over-segmentation seen in Cellpose.

One challenge to using the distance field as the basis to our approach is that traditional distance field algorithms like FMM (Fast Marching Method) are sensitive to boundary pixilation^33^, causing artifacts in the flow field that extend deep into the cell. These artifacts are sensitive to pixel-scale changes at the cell perimeter, which we reasoned would interfere with the training process. To solve this, we developed an alternative approach based on FIM (Fast Iterative Method) that produces smooth distance fields for arbitrary cell shapes and sizes (Fig. 3a, and see Methods)^34^. The corresponding flow field is relatively insensitive to boundary features at points removed from the cell boundary, a critical property for robust and generalized prediction by the Cellpose network.

**Fig. 3.**
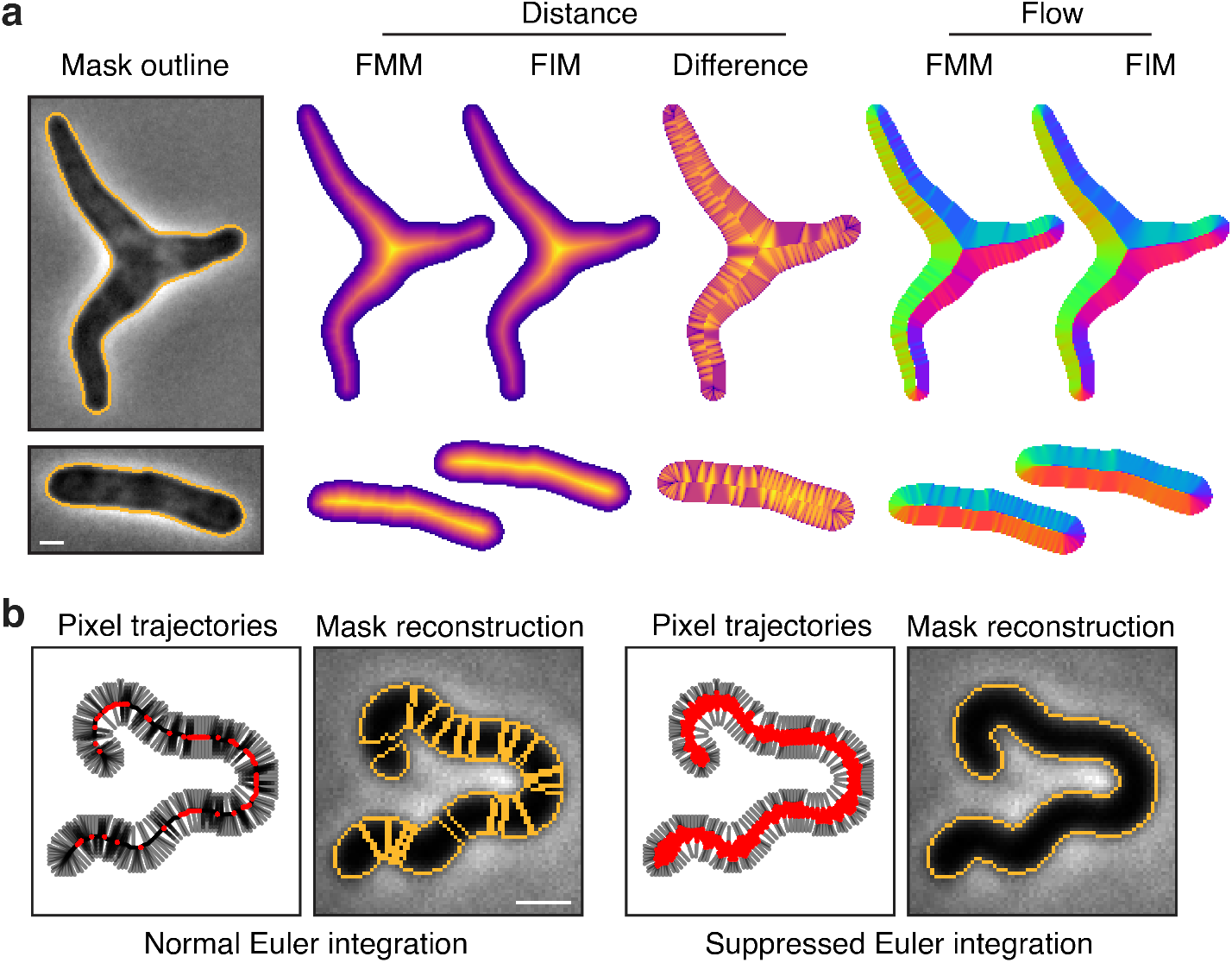
Core innovations of Omnipose. (**a**) Comparison of distance field algorithms and corresponding flow fields. Fast Marching Method (FMM) produces ridges in the distance field resulting from pixelation on the cell mask boundary. Our smooth FIM algorithm minimizes these features. The difference image (FIM – FMM) highlights artifacts in the FMM method. Flow fields are calculated as the normalized gradient of the distance field. Boundary pixelation affects the FMM flow field deep into the cell, regardless of cell size. (**b**) Comparison of mask reconstruction algorithms on a smooth flow field (not shown). Left: boundary pixel trajectories and resulting mask outlines from standard Euler integration. Right: Trajectories and mask outlines under suppressed Euler integration. Red dots indicate the final positions of all cell pixels, not only the boundary pixels for which trajectories are displayed. Bacteria displayed are (a) *Escherichia coli* CS703-1 and (b) and *Helicobacter pylori*, both treated with aztreonam. Scale bars are 1 μm.

The use of the distance field additionally required a unique solution for mask reconstruction. Whereas the pixels in a center-seeking field converge on a point, standard Euler integration under our distance-derived field tends to cluster pixels into multiple thin fragments along the skeleton, causing over-segmentation (Fig. 3b). We solved this with a suppression factor of (*t* + 1)^-1^ in each time step of the Euler integration. This reduces the movement of each pixel after the first step *t* = 0, facilitating initial cell separation while preventing pixels from clustering into a fragmented skeleton formation. The wider distribution resulting from our suppression factor allows pixels to remain connected, thereby generating a single mask for each cell in conjunction with a standard automated pixel clustering algorithm (*e.g*., DBSCAN)^35^.

### Omnipose demonstrates unprecedented segmentation accuracy of bacterial cells

With solutions to the major challenges of cell center-independent segmentation incorporated into Omnipose, we proceeded to benchmark its performance. Remarkably, across the IoU threshold range 0.5-1.0, Omnipose averages a JI >10-fold above that of Cellpose (Fig. 4a). The difference in performance between the algorithms is particularly pronounced within the high IoU range (0.75-1.0), where we observe an average of 170-fold higher JI for Omnipose. At the 0.5-5 μm scale and with a typical microscope configuration, quantitative measurements rely upon IoU values in this range, thus Omnipose is uniquely suited for the microscopic analysis of bacterial cells.

**Fig. 4.**
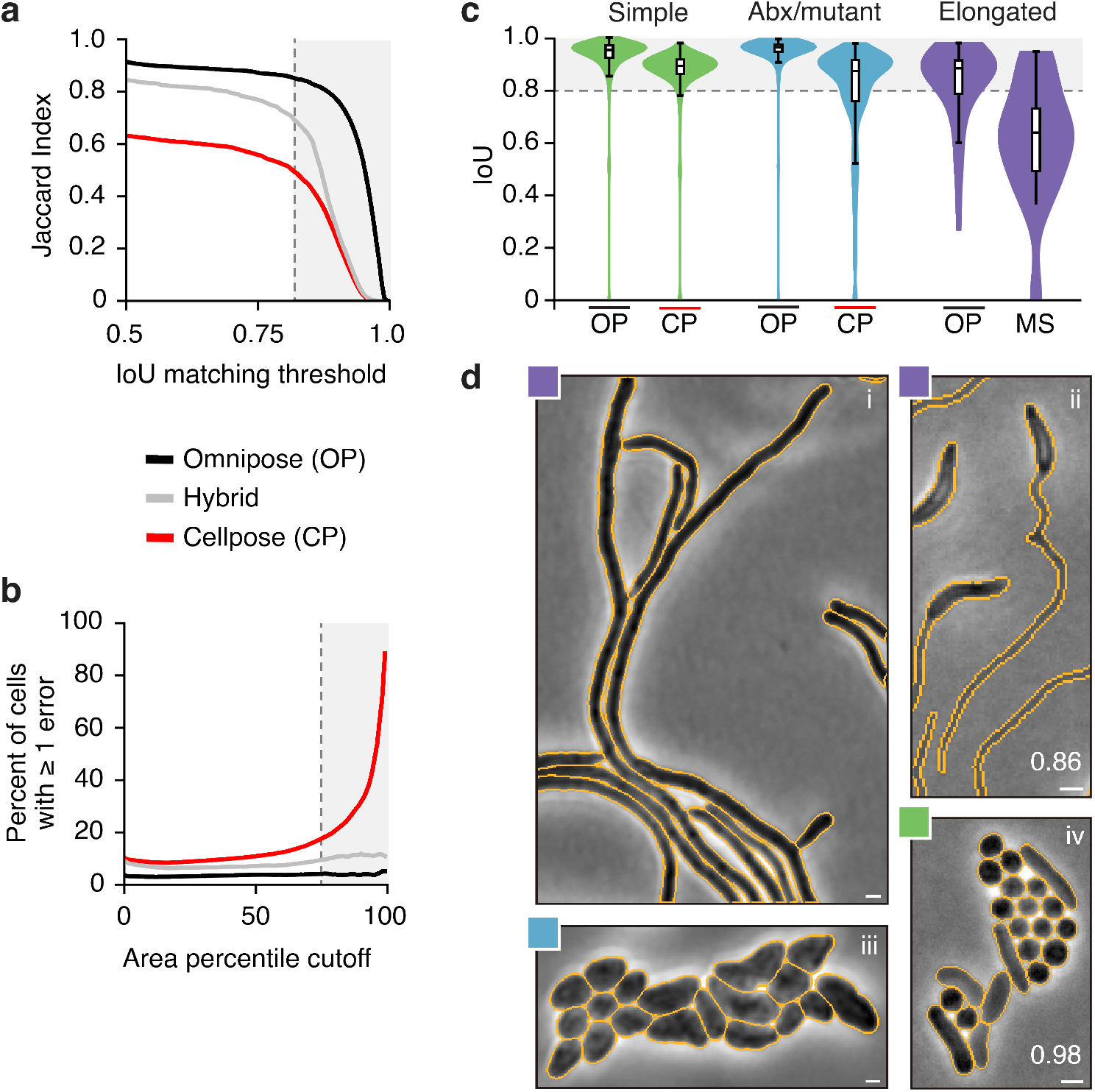
Omnipose outperforms Cellpose. (**a**) Overall performance measured by Jaccard Index (JI). The hybrid method (gray) is a variant of Cellpose that uses the original center-seeking flow output and the mask reconstruction of Omnipose. Gray box represents IoU ≥ 0.8. (**b**) Quantification of segmentation performance by cell size. The percent of cells with at least one segmentation error is computed for cells in each area percentile group from 1 to 100. Gray box represents the top quartile. (**c**) Omnipose IoU distribution on our dataset compared to the next highest performing algorithm in each of three cell categories. (**d**) Example micrographs and Omnipose segmentation. Mean matched IoU values shown. Bacteria displayed are (i) *Streptomyces pristinaespiralis, (ii) Caulobacter crescentus* grown in HIGG media, (*iii*) *Shigella flexneri* treated with A22, (*iv*) mix *of Pseudomonas aeruginosa, Staphylococcus aureus, Vibrio cholerae*, and *Bacillus subtilis*. Scale bars are 1 μm.

To dissect the contributions of the individual Omnipose innovations to the overall performance of the algorithm, we isolated the mask reconstruction component of Omnipose and applied it to the Cellpose network output. This augmentation of Cellpose modestly improved its performance to a roughly equivalent extent across all IoU thresholds (Fig. 4a). Based on this, we attribute the remaining gains in performance by Omnipose to its unique network outputs and our improvements to the Cellpose training framework (see Methods).

Our analyses illuminated critical flaws in prior DNN-based approaches for the segmentation of elongated cells, effectively preventing these algorithms from generalizable application to bacteria (Fig. 1). To determine whether Omnipose overcomes this limitation, we evaluated its performance as a function of cell area. Cell area serves as a convenient proxy for cell length in our dataset, which is composed of both branched and unbranched elongated cells. Whereas the Cellpose cell error rate remains above 9% and increases exponentially with cell size, Omnipose displays a consistent error rate that remains below 4% for all percentiles (Fig. 4b). Thus, Omnipose performance is independent of cell size and shape, including those cells with complex, extended morphologies (Fig. 4c,d).

### Omnipose permits sensitive detection of cellular intoxication

Our laboratory recently described an interbacterial type VI secretion system-delivered toxin produced by *Serratia proteamaculans*, Tre1^36^. We showed that this toxin acts by ADP-ribosylating the essential cell division factor FtsZ; however, we were unable to robustly evaluate the consequences of Tre1 intoxication on target cell morphology owing to segmentation challenges. Here we asked whether Omnipose could permit straightforward and sensitive detection of intoxication by Tre1. To this end, we incubated *S. proteamaculans* wild-type or a control strain expressing inactive Tre1 (*tre1^E415Q^*) with target *E. coli* cells and imaged these mixtures after a fixed period of 20 hours. Owing to the unique capabilities of Omnipose, we were able to include dense fields of view, incorporating >300,000 cells in our analysis.

Among the cells identified by Omnipose, we found a small proportion were elongated and much larger than typical bacteria (Fig. 5a,b and Extended Data Fig. 6a). These cells were only detected in mixtures containing active Tre1, and the apparent failure of the cells to septate is consistent with the known FtsZ-inhibitory activity of the toxin. The *S. proteamaculans* strain background we employed in this work expresses the green fluorescent protein. Corresponding fluorescence images allowed us to unambiguously assign the enlarged cell population to *E. coli* (Fig. 5c). Next, we subjected the same images to cell segmentation with StarDist, Cellpose, and MiSiC, the three top-performing algorithms in our initial survey. Each of these algorithms fail to identify this population of cells to high precision (Fig. 5d,e). Close inspection reveals three distinct modes of failure (Fig. 5e and Extended Data Fig. 6b). In the case of StarDist, elongated (non-star-convex) cells are split into multiple star-convex subsets that do not span the entire cell. Cellpose detects entire elongated cells, but it breaks them up into a multitude of smaller masks. Conversely, MiSiC detects all cells but fails to properly separate them, thereby exaggerating the area measurement in many cases. Taken together, these data illustrate how the enhanced cell segmentation performance of Omnipose can facilitate unique insights into microbiological systems.

**Fig. 5.**
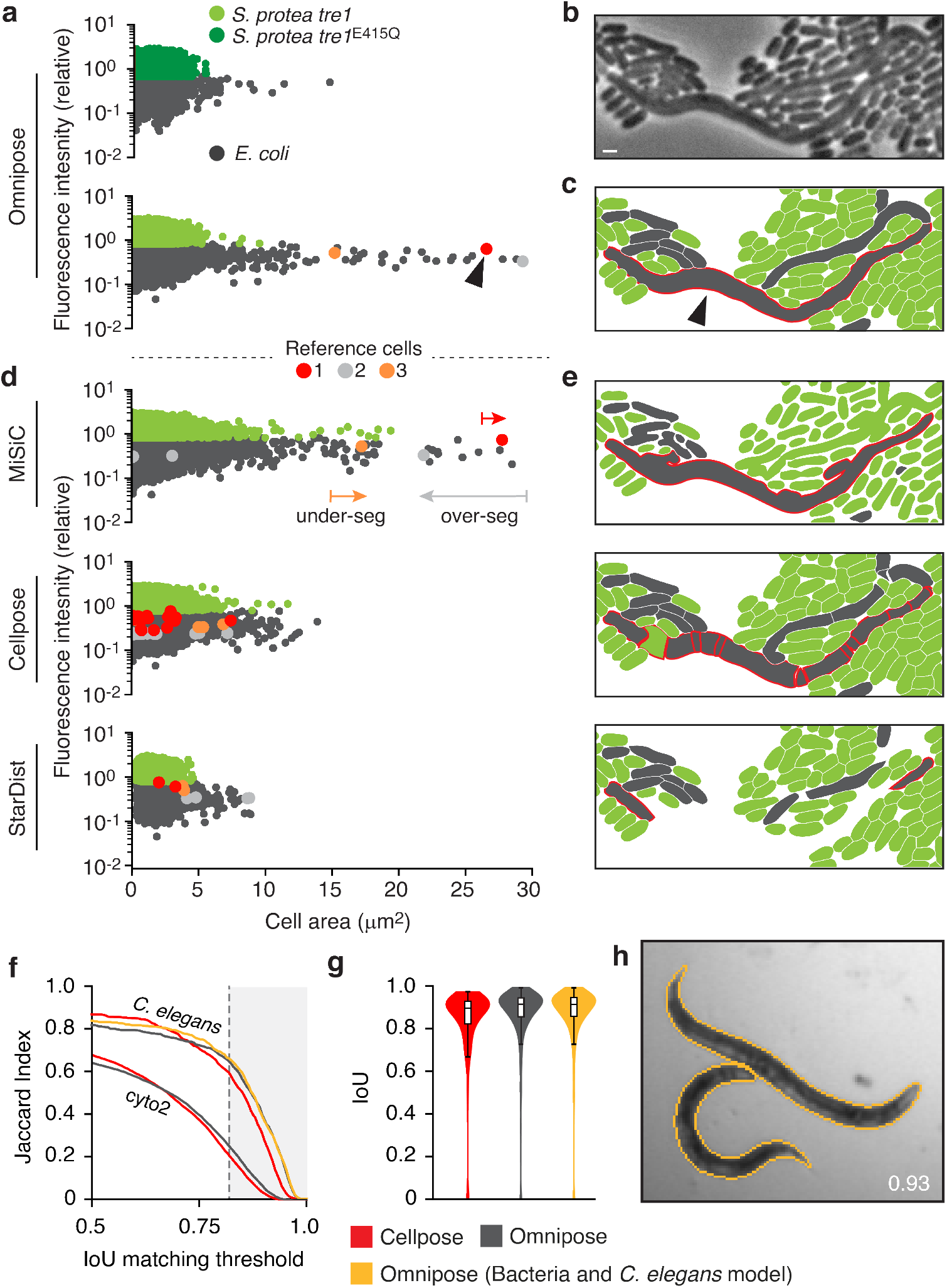
Omnipose can be applied to the study of bacterial and non-bacterial systems. (**a**) Fluorescence/area population profile according to Omnipose segmentation in control and experimental conditions. K-means clustering on GFP fluorescence distinguishes *S. proteamaculans tre1/tre1^E415Q^* (light/dark green markers) from *E. coli* (gray markers). (**b**) Example of extreme filamentation of *E. coli* in response to active Tre1. (**c**) Omnipose accurately segments all cells in the image. Largest cell indicated with black arrow. (**d**) MiSiC predicts large cell masks over both species. Cellpose and StarDist fail to predict any cells above 15μm^2^. (**e**) Example segmentation results highlighting typical errors encountered with MiSiC (under-segmentation), Cellpose (over-segmentation), and StarDist (incomplete masks). Mask mergers cause some *E. coli* to be misclassified as *S. proteamaculans*. Scale bar is 1 μm. (**f**) Performance of Omnipose and Cellpose on cyto2 and *C. elegans* datasets. Results for Omnipose trained on *C. elegans* (grey) or *C. elegans* and bacterial data (yellow) are shown. (**g**) IoU distribution for the masks predicted by each method on our *C.elegans* dataset. (**h**) Example segmentation of *C. elegans* in the BBBC010 dataset.

### Omnipose exhibits strong performance in non-bacterial segmentation tasks

We have shown that the features we developed within Omnipose improved phase-contrast bacterial segmentation performance beyond that of Cellpose. However, it is possible that these features could hinder performance relative to Cellpose in other imaging modalities or in segmentation tasks involving, for example, eukaryotic cells, whole organisms, and cell-like objects. To test this, we trained Omnipose on the cyto2 dataset, a large collection of images and corresponding ground-truth annotations submitted by users that expands upon the original cyto dataset of Cellpose^12,30^. We found that Omnipose offers a modest improvement in performance relative to Cellpose on this dataset (Fig. 5f and Extended Data Fig. 7a). Moreover, Omnipose achieved this performance boost without compromising the segmentation rate (~1 image per second).

Encouraged by the strong performance of Omnipose on the cyto2 dataset, we next sought to investigate its potential utility in a field far removed from microbiology. The nematode *Caenorhabditis elegans* is a widely studied model organism with an overall morphology grossly similar to elongated bacteria^37^. At just one millimeter in length, intact *C. elegans* are often analyzed by microscopy in order to measure phenotypes; therefore, there is significant interest in methods for their accurate segmentation to enable tracking^38^. We obtained, annotated and trained Omnipose on two publicly available microscopy datasets composed of *C. elegans* images: time-lapse frames from the Open Worm Movement database^39^ and frames containing fields of assorted live or dead *C. elegans* from the BBBC010 dataset^40^. These images contain debris and are of heterogenous quality, yet 82% of masks predicted by Omnipose match or exceed the 0.8 IoU threshold (Fig. 5g,h). Taken together with our findings on cyto2, we conclude that Omnipose inherits and offers improvement upon the broad applicability of Cellpose.

## Discussion

Confronted with the importance of segmentation accuracy to the success of work within our own laboratory, we were motivated to characterize the performance of several existing cell segmentation algorithms. Recent developments in deep learning have greatly improved these algorithms; however, significant challenges remain^1,30^. Although isolated cells without cell-to-cell contact can be segmented with high precision by any of the packages we tested, segmentation becomes significantly more challenging when cells form microcolonies, adopt irregular morphologies, or when fields are composed of cells with multiple shapes and sizes. Such difficulties are compounded in time-lapse studies, where the significance of segmentation errors often grows exponentially with time. Experimental design can help mitigate certain segmentation challenges; however, the recent emphasis on non-model organisms and microbial communities renders this an increasingly undesirable solution^41^.

This work provides the most comprehensive side-by-side quantitative comparison of cell segmentation algorithm performance to-date. As expected, machine-learning-based approaches outperform others, yet insights into general image segmentation strategies can be gained from each of the methods we examined. Two of the six algorithms we tested utilize traditional image thresholding and watershed segmentation: Morphometrics and SuperSegger^13,23^. Each program tends to under-segment adjacent cells and over-segment large cells, behaviors previously linked to thresholding and watershed processes, respectively^1,42^. Given that SuperSegger was motivated at least in-part to mitigate these issues, we postulate that traditional image segmentation approaches are ultimately limited to specialized imaging scenarios. Although we classify MiSiC as a DNN-based approach, this algorithm also relies on thresholding and watershed segmentation to generate cell masks from its network output^15^. The network output of MiSiC is more uniform than unfiltered phase contrast images, yet this pre-processing does not fully abrogate the typical errors of thresholding and watershed segmentation. We therefore conclude that, even when combined with neural networks as seen in MiSiC, thresholding and watershed cannot be effectively used for general cell segmentation tasks.

A successful DNN-based algorithm is composed of a robust, consistent neural network output, and an appropriate mask reconstruction process based on this output. In the case of Mask R-CNN, bounding boxes for each cell are predicted along with a probability field that localizes a cell within its bounding box^43^. Masks are generated by iterating over each box and thresholding the probability field. Despite the widespread adoption of Mask R-CNN, we found this algorithm did not perform exceptionally well in our study. Our results suggest that this is due to dense cell fields with overlapping bounding boxes, a feature known to corrupt the training process and produce poor network outputs for Mask R-CNN^44^. By contrast, the StarDist network makes robust predictions, but it fails to assemble accurate cell masks because the cells in our dataset are not well approximated by star-convex polygons^26^. The errors we encountered with Cellpose can be attributed to both neural network output and mask reconstruction. In Omnipose, we specifically addressed these two issues via the distance field and suppressed Euler integration, respectively, yielding a remarkably precise and generalizable image segmentation tool. Omnipose effectively leverages the strongest features of several of the DNN approaches we tested, namely the distance field of StarDist, the boundary field of MiSiC, and the mask reconstruction framework of Cellpose.

We have designed Omnipose for use by typical research laboratories and we have made its source code and training data publicly available. For images of bacteria under phase contrast, researchers will not need to provide new ground truth data or retrain the model. In this study, we emphasized morphological diversity, but we further accounted for differences in optical features between bacterial strains, slide preparation techniques, and microscope configurations. For example, the images in our ground-truth dataset originate from four different researchers using distinct microscopes, objectives, sensors, illumination sources, and acquisition settings. We further introduced extensive image augmentations that simulate variations in image intensity, noise, gamma, clipping, and magnification. Lastly, bacterial strains exhibit a wide range of intrinsic contrast and internal structure, often exacerbated by antibiotic treatment or revealed by dense cell packing. Internal structure can cause over-segmentation, so we included many cells with this characteristic in our dataset.

We anticipate that the unprecedented performance of Omnipose may permit access to information from microscopy images that was previously inaccessible. For instance, images deriving from natural microbial communities could be accurately characterized with regard to internal structure, autofluorescence, and morphology at the single-cell level. This data could be used to estimate diversity, a novel methodology that would complement existing sequencing-based metrics^45^. It is worth noting that phenotypic diversity often exceeds genetic diversity^46^; therefore, even in a relatively homogeneous collection of organisms, precise segmentation could allow classes representing distinct states to be identified. A microscopy-based approach also offers the opportunity to characterize spatial relationships between cells, information that is exceptionally difficult to recover in most biomolecular assays.

## Methods

### Phase contrast and fluorescence microscopy

In-house imaging was performed on a Nikon Eclipse Ti-E wide-field epi-fluorescence microscope, equipped with a sCMOS camera (Hamamatsu) and X-cite LED for fluorescence imaging. We imaged through 60X and 100X 1.4 NA oil-immersion PH3 objectives. The microscope was controlled by NIS-Elements. Cell samples were spotted on a 3% (w/v) agarose pad placed on a microscope slide. The microscope chamber was heated to 30°C or 37°C when needed for time-lapse experiments.

Several images in our dataset were taken by two other laboratories using three distinct microscope/camera configurations. The Brun lab provided images of *C. crescentus* acquired on a Nikon Ti-E microscope equipped with a Photometrics Prime 95B sCMOS camera. Images were captured through a 60X Plan Apo λ 100X 1.45 NA oil Ph3 DM objective. The Wiggins lab provided *E. coli* and *A. baylyi* time lapses from both a Nikon Ti-E microscope as well as a custom-built tabletop microscope, both described in previous studies^47,48^.

### *C. elegans* data preparation

We obtained a 1000-frame time lapse of *C. elegans* from the Wormpose^38^ GitHub (https://github.com/iteal/wormpose_data) adapted from the Open Worm Movement database^39^, which is inaccessible at the time of writing. We also utilized the Broad Bioimage Benchmarking Collection set BBBC010^40^ (https://bbbc.broadinstitute.org/c-elegans-livedead-assay-0), a set of 100 images containing live and dead *C. elegans*. These images were manually cropped to select regions of each image without *C. elegans* overlaps. For both of these datasets, images were initially segmented with Omnipose to select foreground, automatically cropped to select individual *C. elegans* or clusters of *C. elegans*, and then packed into ensemble images for efficient annotation, training, and testing following the same procedures described below for our bacterial datasets.

### Bacterial sample preparation

To image antibiotic-induced phenotypes, cells were grown without antibiotics overnight in LB, back-diluted, and spotted on agarose pads with 50μg/mL A22 or 10μg/mL cephalexin. Time lapses were captured of *E. coli* DH5α and *S. flexneri* M90T growing on these pads. *E. coli* CS703-1 was back-diluted into LB containing 1μg/mL aztreonam and spotted onto a pad without antibiotics^49^. Cells constitutively expressed GFP to visualize cell boundaries.

*H. pylori* LSH100 grown with and without Aztreonam was provided by the Salama lab^50,51^. Samples were fixed and stained with Alexaflour 488 to visualize the cell membrane. Images were taken on LB pads. The typical technique of allowing the spot to dry on the pad caused cells to curl up on themselves, so our images were taken by placing the cover slip on the pad immediately after spotting and applying pressure to force out excess media.

*C. crescentus* was cultivated and imaged by the Brun lab^52,53^. Cells were grown in PYE, washed twice in water prior to 1:20 dilution in Hutner base-imidazole-buffered-glucose-glutamate (HIGG media) and grown at 26°C for 72h. Cells were spotted on a 1% agarose PYE pads prior to imaging.

*S. pristinaespiralis* NRRL 2958 was grown using the following media recipe: Yeast extract 4g/L, Malt extract 10g/L, Dextrose 4g/L, Agar 20g/L. This media was used to first culture the bacteria in liquid overnight and then on a pad under the microscope. This strain forms aggregates in liquid media, so these aggregates were allowed to grow for several hours on a slide in the heated microscope chamber until we could see individual filaments extending from the aggregates. Fields of view were selected and cropped to exclude cell overlaps. Autofluorescence was captured to aid in manual segmentation.

Mixtures of *S. proteamaculans* attTn7::Km-gfp *tre1* or *tre1^E415Q^* and *E. coli* were spotted on a PBS pad to prevent further growth. Phase-contrast images of the cells were acquired before and after a 20hr competition on a high-salt LB plate. Fluorescence images in the GFP channel were also acquired to distinguish *S. proteamaculans* from unlabeled *E. coli*.

All other individual strains in Table S1 were grown overnight, diluted 1:100 into fresh LB media, and grown for 1-3 hours before imaging. Mixtures were made by combining back-diluted cells roughly 1:1 by OD_600_.

### Manual image annotation

Manual annotation began with loading images into MATLAB, normalizing the channels, registering the fluorescence channel(s) to brightfield (when applicable), and producing boundary-enhanced versions of brightfield and fluorescence. Where possible, fluorescence data was primarily used to define cell boundaries (not available in the *C. elegans* dataset acquired online). In addition to a blank channel to store manual labels, all processed phase and fluorescence images were then automatically loaded as layers into an Adobe Photoshop document. We used 4-6 unique colors and the Pencil tool (for pixel-level accuracy and no blending) to manually define object masks. Due to the 4-color theorem^54^, this limited palette was sufficient to clearly distinguish individual object instances from each other during annotation. This color simplification is not found in any segmentation GUI, and it enabled faster manual annotation by reducing the need to select new colors. It also eliminated the confusion caused by the use of similar but distinct colors in adjacent regions, which we suspect is the principal cause for the misplaced mask pixels that we observed in other datasets (*e.g*., cyto2).

The cell label layer was then exported as a PNG from Photoshop, read back into MATLAB, and converted from the repeating N-color labels to a standard 16-bit integer label matrix, where each object is assigned a unique integer from 1 to the number of cells (background is 0). Because integer labels cannot be interpolated, we then performed a non-rigid image registration of the brightfield channel to the binary label mask to achieve better brightfield correlation to ground truth masks. All images in our ground-truth dataset have been registered in this manner.

### Choosing Segmentation algorithms

Three main factors contributed to the choice of algorithms highlighted in this study: (*i*) specificity to bacterial phase contrast images, (*ii*) success and community adoption, especially for bioimage segmentation, and (*iii*) feasibility of installation, training, and use. SuperSegger, Morphometrics, and MiSiC were selected because they specifically targeted the problem of bacterial phase contrast segmentation^15,23,55^. Packages such as BactMAP, BacStalk, Cellprofiler, CellShape, ColiCoords, Cytokit, MicroAnalyzer, MicrobeJ, Oufti, and Schnitzcells incorporate limited novel segmentation solutions and instead aim to provide tools for single-cell analysis such as lineage tracing and protein tracking^8,9,14,18,20,25,56,58^. Furthermore, the segmentation that these programs perform depends broadly on thresholding and watershed techniques; therefore, Morphometrics is a reasonable proxy for their segmentation capabilities. We were unable to locate code or training data for BASCA^11^. Ilastik is a popular interactive machine-learning tool for bioimage segmentation, but training it using a manual interface was not feasible on a large and diverse dataset such as our own^21^. Among DNN approaches, Mask R-CNN was selected because it is a popular architecture for handling typical image segmentation tasks. It was also used in the segmentation and tracking package Usiigaci^24^. U-Net architectures have been implemented in a number of algorithms, including DeLTA, PlantSeg, MiSiC, StarDist, and Cellpose^12,15,17,22,26^. DeLTA was not included in this study because it operates similarly to MiSiC and was designed specifically for mother machine microfluidics analysis. DeLTA 2.0 was recently released to additionally segment confluent cell growth on agarose pads, but it remains quite similar to MiSiC in implementation^59^. PlantSeg could, in principle, be trained on bacterial micrographs, but we determined that its edge-focused design meant to segment bright plant cell wall features would not offer any advancements over the remaining U-Net methods that we tested.

### Training and tuning segmentation algorithms

All segmentation algorithms have tunable parameters to optimize performance on a given dataset. These include pre-processing such as image rescaling (often to put cells into a particular pixel diameter range), contrast adjustment, smoothing, and noise addition. Morphometrics and SuperSegger were manually tuned to give the best results on our benchmarking dataset. The neural network component of SuperSegger was not retrained on our data, as this is a heavily manual process involving toggling watershed lines on numerous segmentation examples. DNN-based algorithms are automatically trained using our dataset, and the scripts we used to do so are available in our GitHub repository. We adapted our data for MiSiC by transforming our instance labels into interior and boundary masks. Training documentation for MiSiC is not published. Training and evaluation parameters for MiSiC were tuned according to correspondence with the MiSiC authors. Cellpose and StarDist were trained with the default parameters provided in their documentation. StarDist has an additional tool to optimize image pre-processing parameters on our dataset, which we utilized.

### Evaluating segmentation algorithms

All algorithms were evaluated on our benchmarking dataset with manually or automatically optimized parameters. We provide both the raw segmentation results for all test images by each tested algorithm as well as the models and model-training scripts required to reproduce our results. Before evaluating IoU or JI, small masks at image boundaries were removed for both the ground-truth and predicted masks. IoU and JI are calculated on a per-image basis and, where shown, are averaged with equal weighting over the image set or field of view.

Our new metric, the number of segmentation errors per cell, was calculated by first measuring the fraction of each predicted cell that overlaps with each ground truth cell. A predicted cell is assigned to a ground-truth cell if the overlap ratio is ≥ 0.75, meaning that at least three quarters of the predicted cell lies within the ground-truth cell. If several predicted cells are matched to a ground-truth cell, the number of surplus matches is taken to be the number of segmentation errors. If no cells are matched to a ground-truth cell, then the error is taken to be 1.

### Leveraging Omnipose to accelerate manual annotation

Omnipose was periodically trained on our growing dataset to make initial cell labels. These were converted into an N-color representation and loaded into Photoshop for manual correction. A subset of our cytosol GFP channels were sufficient for training Omnipose to segment based on fluorescence, and the resulting trained model enabled higher-quality initial cell labels for GFP-expressing samples than could be achieved from intermediate phase contrast models (*e.g., V. cholerae*).

### Defining the Omnipose prediction classes

Omnipose predicts four classes: two flow components, the distance field, and a boundary field. Our distance field is found by solving the eikonal equation

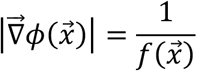

where *f* represents the speed at a point 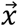. The Godunov upwind discretization of the eikonal equation is

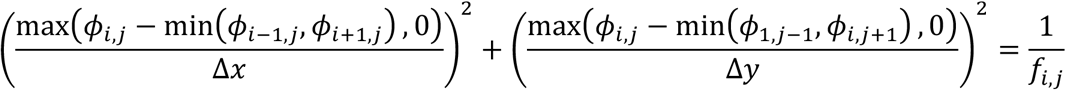

Our solution to this equation is based on the Improved FIM Algorithm 1.1 of^34^, as follows. Our key contribution to this algorithm is the addition of ordinal sampling to boost both convergence and smoothness of the final distance field.

*2D update function for ϕ_i,f_ on a cartesian grid*

1. Find neighboring points for cardinal axes (Δ*x* = Δ*y* = *δ*):

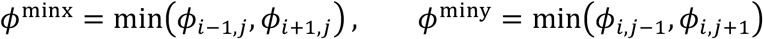
2. Find neighboring points for ordinal axes 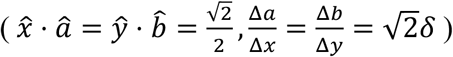:

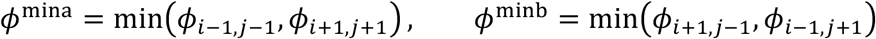
3. Calculate update along cardinal axes:

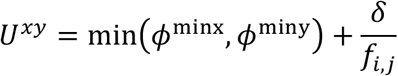

**else**:

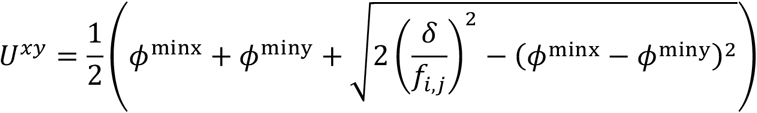
4. Calculate update along ordinal axes:

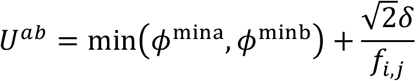

**else**:

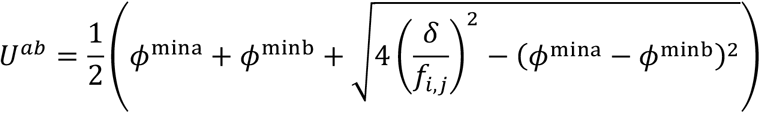
5. Update with geometric mean:

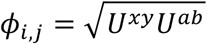

This update rule is repeated until convergence (Extended Data Fig. 5). We take *δ* = *f_i,j_* to obtain the signed distance field used in Omnipose. The flow field components are defined by the normalized gradient of this distance field *ϕ*. The boundary field is defined by points satisfying 0 < *ϕ* < 1. For network prediction, the boundary map is converted to the logits (inverse sigmoid) representation, such that points in the range [0,1] are mapped to [-5,5]. For consistent value ranges across prediction classes, the flow components are multiplied by 5 and all background values of the distance field (*ϕ* = 0) are set to −5.

### Omnipose network architecture

The DNN used for Omnipose is a minor modification of that used in Cellpose: a U-net architecture with two residual blocks per scale, each with two convolutional layers^12^. Omnipose introduces a dropout layer before the densely connected layer^60^, which we incorporated into the shared Cellpose and Omnipose architecture moving forward. However, Cellpose models utilized in this study are trained without dropout.

### Rescaling flow field by divergence

During training, the ground truth data is augmented by a random affine transformation. The original implementation, and the one which yields the best results, linearly interpolates the transformed field. This reduces the magnitude of the otherwise normalized field in regions of divergence, *i.e*., at boundaries and skeletons. A renormalized field (obtained either from the transformed field or as the normalized gradient of the transformed heat distribution) often has artifacts at cell boundaries and skeletons, so the interpolated field effectively reduces the influence of these artifacts on training. We reason that this feature explains the superior performance of interpolated field training over renormalized fields, despite the latter being the nominal goal of the algorithm.

Pixels at cell boundaries, however, consequently do not move far (less than 1px) under Euler integration due to the low magnitude of the predicted field at cell boundaries. Our solution in Omnipose is to rescale the flow field by the magnitude of the divergence. The divergence is most positive at the cell boundaries (where pixels need to move) and most negative at cell skeletons (where pixels need to stop). We therefore rescale the divergence from 0 to 1 and multiply the normalized flow field by this new magnitude map. This forces boundary pixels of neighboring cells to quickly diverge and allow for accurate pixel clustering to obtain the final segmentation.

### Novel diameter metric

The size models of Cellpose are trained to estimate the average cell ‘diameter’, taken to be the diameter of the circle of equivalent area:

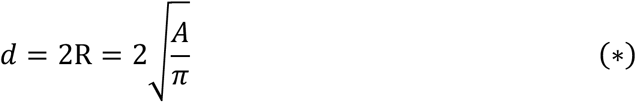

This metric as a basis for rescaling is problematic when cells are growing in length but not width (Extended Data Fig. 7). Log-phase bacterial cell area grows exponentially with time, and so too does the scale factor, eventually resulting in a rescaled image that is too small for Cellpose to segment.

The average of the distance field, however, does not change for filamentous bacteria, as the width – and therefore the distance to the closest boundary – remains constant. To define a formula consistent with the previous definition in the case of a circular cell, we consider mean of the distance field over the cell:

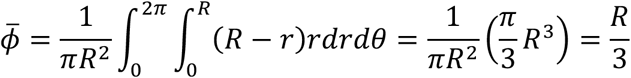

This allows us to define a new ‘effective diameter’ as

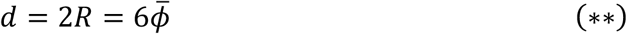

Aside from agreeing with the previous scaling method (*) for round morphologies, this definition exhibits excellent consistency across time (Extended Data Fig. 7). This consistency is also critical for training on datasets with wide distributions in cell areas that require rescaling, such as the Cellpose datasets. Finally, the raw distance field output of Omnipose can directly be used directly in (**) to estimate average cell diameter, which is used in our code to automatically toggle on features that improve mask reconstruction performance for small cells.

### Gamma augmentation

To make the network robust against changes in exposure/contrast, the training images are now raised to a random power (gamma) between 0.5 and 1.25, simulating the varying levels of contrast that are observed experimentally with different light sources, objectives, and exposure times.

### Alleviating class imbalance

Class imbalance remains a challenge in many machine learning applications^61^. In our dataset, foreground pixels (cells) take up anywhere from 1 to 75 percent of a given training image, with the rest being background pixels that the network must only learn to ignore (*i.e*., assign a constant output of −5 for distance and boundary logits). We implemented several changes to the loss function to emphasize foreground objects, including weighting by the distance field and averaging some loss terms only over foreground pixels. Our training augmentation function also attempts many random crop and resizing passes until a field of view with foreground pixels is selected (this may take several attempts for sparse images, but adds very little time to training).

### Image normalization

To manage different image exposure levels, Cellpose automatically rescales images such that pixels in the 1st percentile of intensity are set to 0 and those in the 99th percentile are sent to 1. This percentile rescaling is preferred over blind min-max rescaling because bubbles or glass can cause small bright spots in the image. However, we found that images containing single cells (low intensity) in a wide field of media (high intensity) would become badly clipped due to the foreground-background class imbalance. To solve this, we changed the percentile range from 0.01 to 99.99.

## Data availability

Ground truth images and labels generated for this study are hosted on the Cellpose website (http://www.cellpose.org/dataset_omnipose) and listed on the Papers With Code database (https://paperswithcode.com/dataset/bpcis).

## Code availability

For install instructions, source code, and all Python and MATLAB scripts generated for this study, see our GitHub repository at https://github.com/kevinjohncutler/omnipose.

## Acknowledgements

The authors wish to thank members of the Mougous and Wiggins laboratories for helpful suggestions, See-Yeun Ting for assistance with experiments, Teresa Lo for assistance with image acquisition, Sophie Sichel and Nina Salama lab for growing, fixing, and staining *H. pylori* samples for in-house imaging, and David Kysela, Maxime Jacq and Yves Brun for providing *C. crescentus* images. This work was supported by the NIH (AI080609 to JDM, GM128191 to PAW, T32GM008268 to KJC). JDM is an HHMI Investigator.

## Author contributions

KJC, PAW and JDM conceived the study. KJC performed experiments, analyzed data, and wrote the code. KJC, PAW, and JDM wrote the manuscript. CS edited the manuscript and assisted in code development.

## Author Information

Correspondence and requests for materials should be addressed to J.D.M. or P.A.W.

---

**Extended Data Fig. 1.**
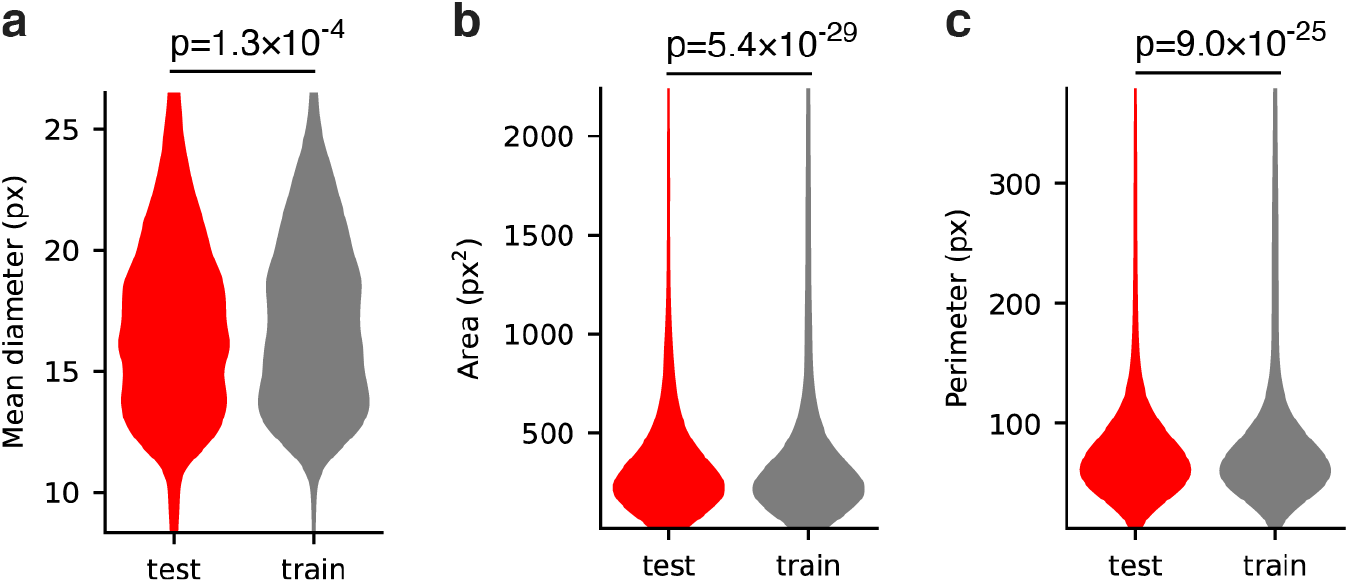
Test dataset is representative of the training dataset. (**a**) Mean diameter, defined in Methods. (**b**) Cell area. (**c**) Cell perimeter. P-values are displayed for the two-sided KS test.

**Extended Data Fig. 2.**
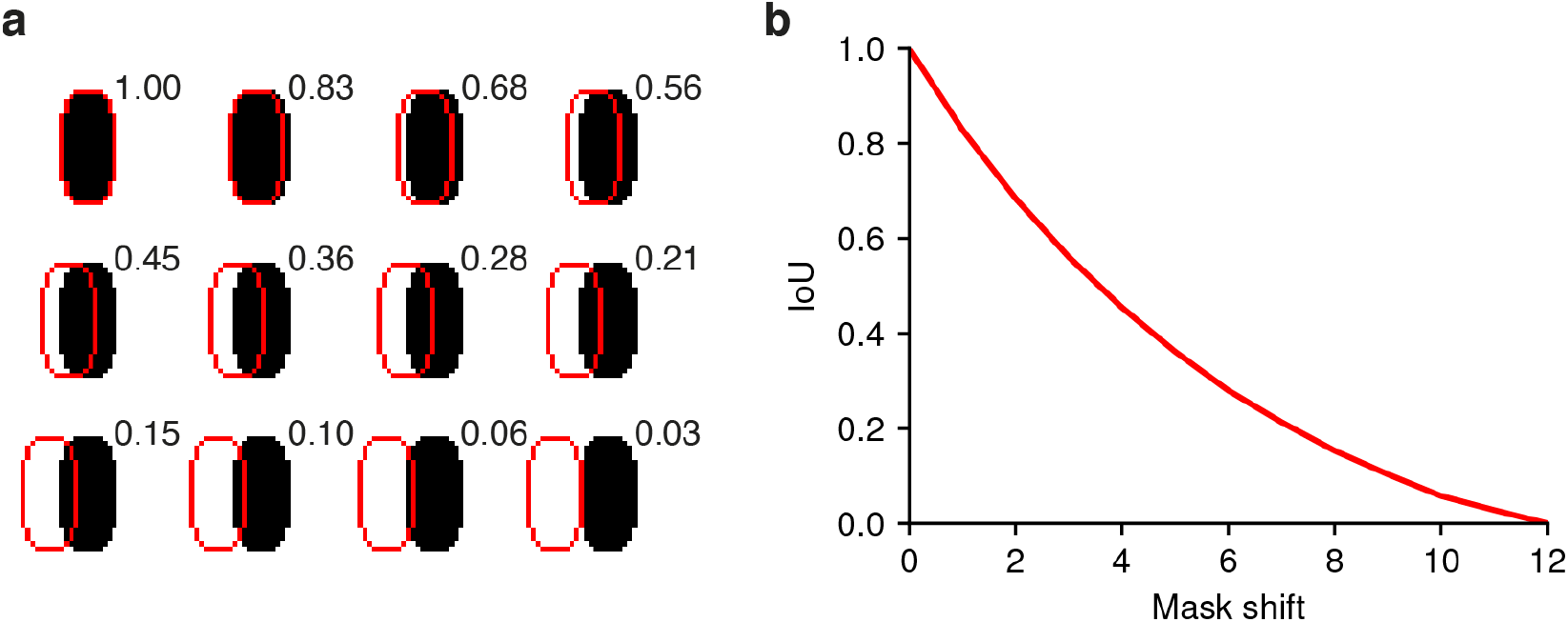
IoU values for synthetic cell of typical size/resolution. (**a**) 0-12 pixel displacement of cell mask (red outline) and corresponding IoU values. (**b**) IoU decreases non-linearly for curved regions such as this synthetic cell.

**Extended Data Fig. 3.**
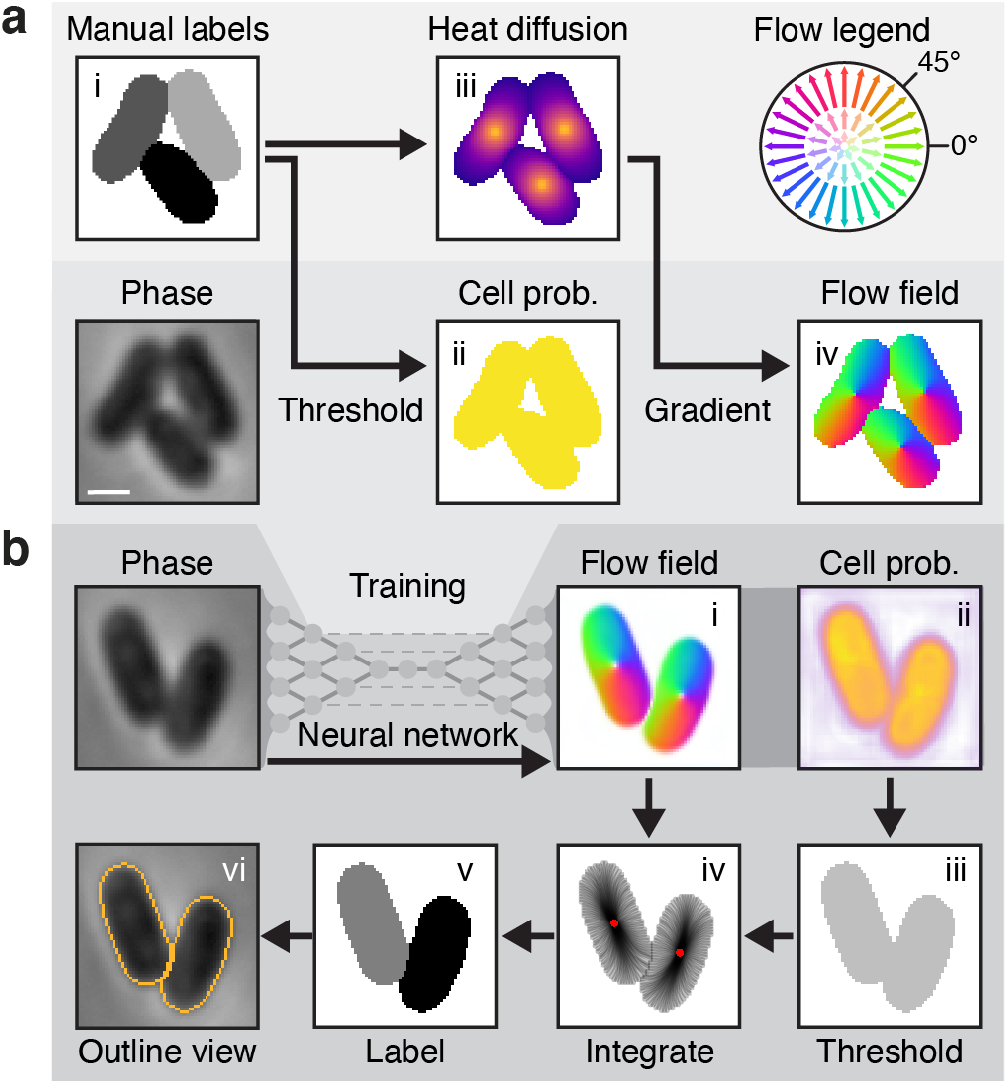
Details of the Cellpose algorithm. (**a**) Stages of the Cellpose training pipeline. Ground truth masks (*i*) are converted to cell probability (*ii*) by binary thresholding and a heat distribution (*iii*) by simulated diffusion from the median pixel coordinate. The flow field (*iv*) is defined by the normalized gradient of (*iii*). Color-magnitude representations of this vector field follow the flow legend diagram. The phase, cell probability, and flow fields are used to train the network. (**b**) Stages of the cellpose prediction pipeline. Phase images are processed by the trained cellpose network into the intermediate flow field and cell probability outputs (*i-ii*). A binary threshold is applied to the probability to identify cell pixels (*iii*). Pixels are Euler-integrated under the flow field until they converge at common points. Boundary pixel trajectories are depicted in *iv*. Each pixel is assigned a unique label corresponding to the center to which it converged (*v*). This segmentation result is commonly depicted in an outline view (*vi*). Bacteria shown are *Escherichia coli*. Scale bar is 1 μm.

**Extended Data Fig. 4.**
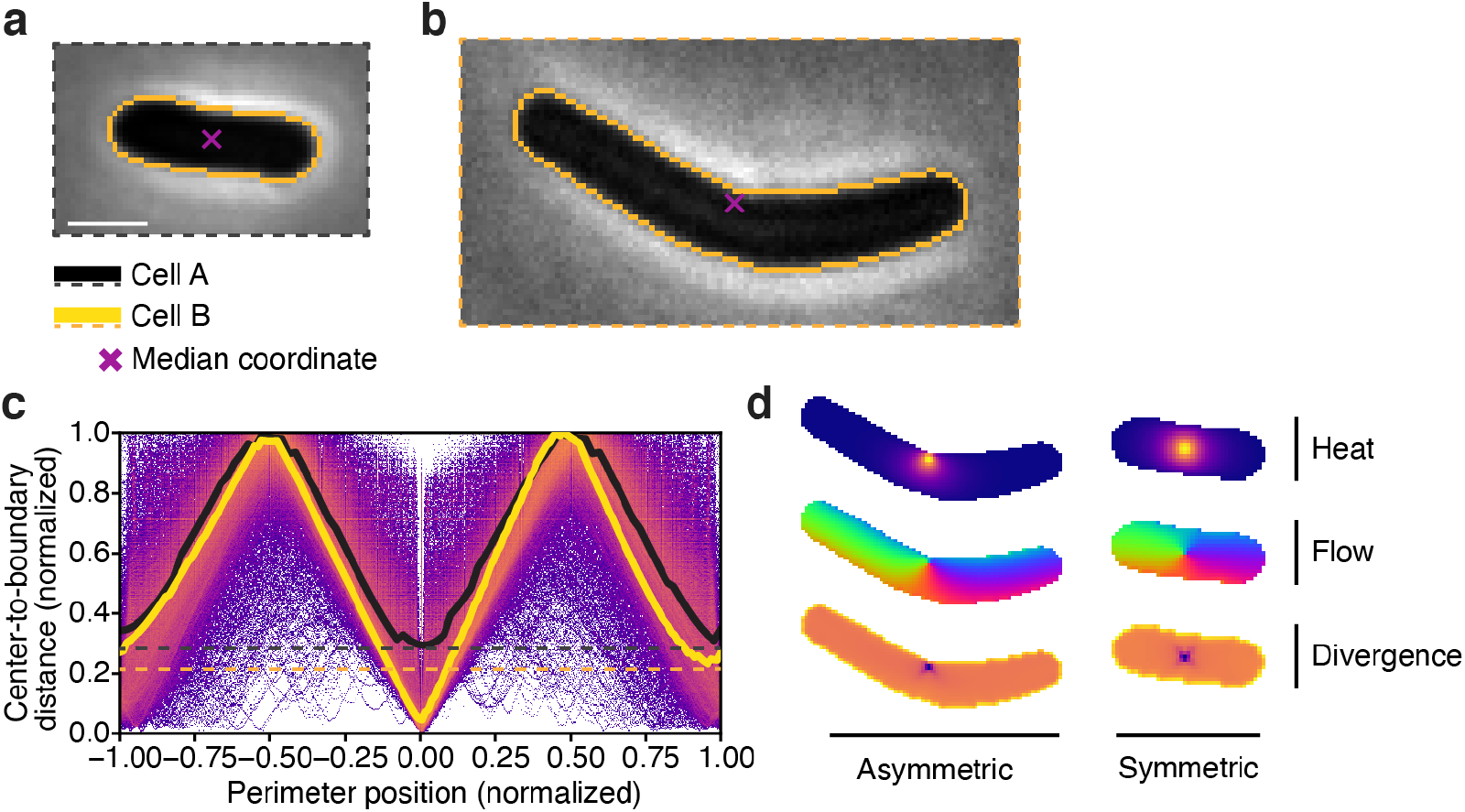
Median coordinates are asymmetrically localized. (**a**) Center-to-boundary distance highlighted for two cells with non-projected median coordinates. Dashed lines indicate the larger of the two minima along the medial axis. (**b**) Rod-shaped *E.coli* with symmetric median coordinate. Symmetry of the center is reflected in A by equal high and low points corresponding to the extremal points along the long and short axes of the cell. (**c**) Curved *B. subtilis* with median coordinate asymmetrically close to the cell boundary. This asymmetry is reflected in A by a secondary minimum above the global minimum corresponding to the diametrically opposing point along the short axis of the cell. (**d**) These centers result in distinct flow fields reflecting the (a)symmetric of the cell center. Bacteria shown are (a) *Escherichia coli* and (b) *Bacillus subtilis*. Scale bar is 1 μm. Images scaled equivalently.

**Extended Data Fig. 5.**
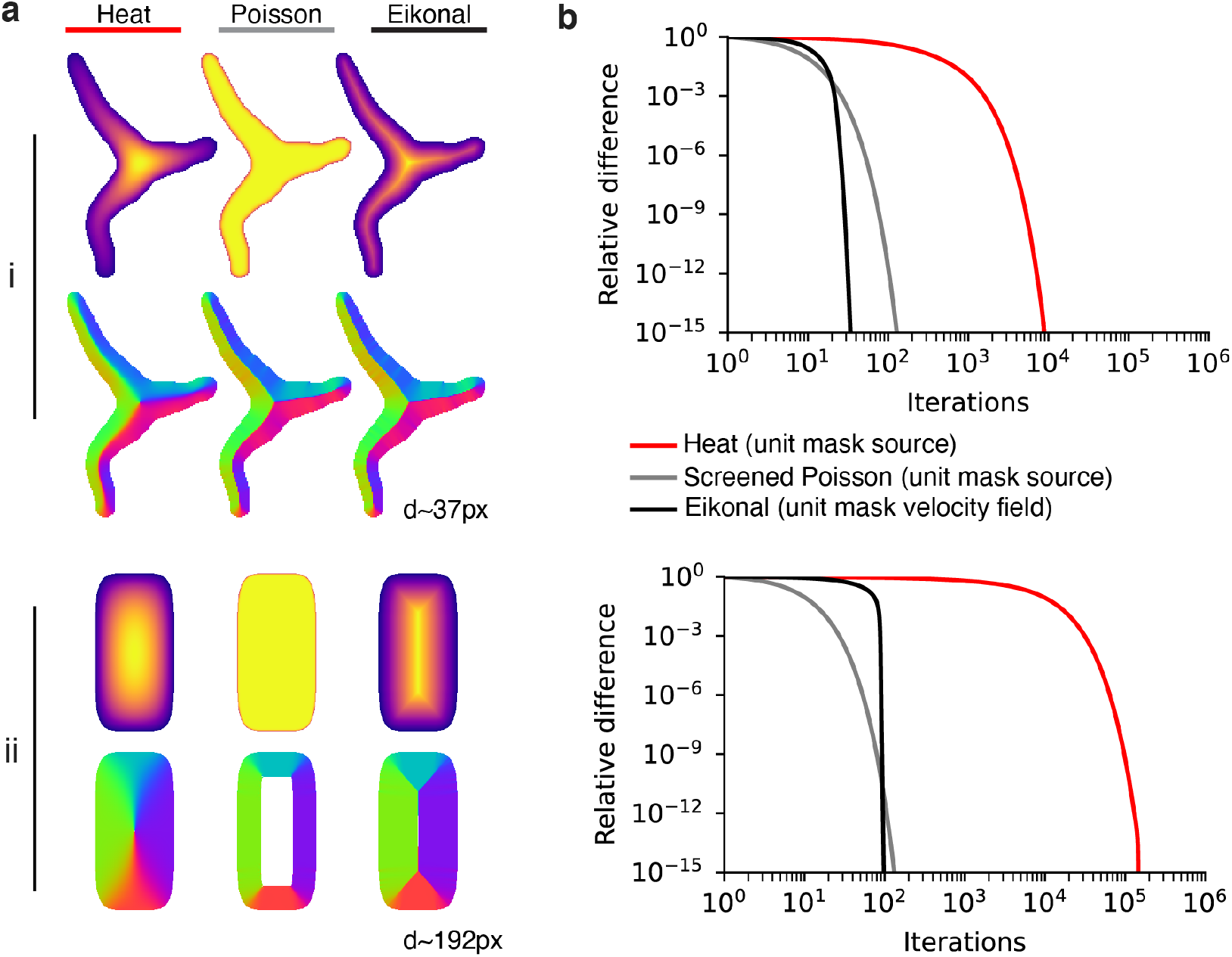
Comparison of three algorithms for computing center-independent flow fields. Each field is defined by a partial differential equation with the mask at the source: time-independent heat equation, the screened Poisson equation, and the Eikonal equation. We solve these equations with iterative relaxation (see Methods). (**a**) Two example cells, the first drawn from our dataset with a mean diameter of 37px and a synthetic rod-shaped cell with a mean diameter of 192px. Cell (*i*) exhibits heat-derived flow components pointing toward the skeleton near boundaries and toward the global cell center at the skeleton. Center-seeking flow components become problematic for mask reconstruction for more complicated cell geometries, namely those with oscillating thickness. The screened Poisson and Eikonal equations produce nearly identical flow fields (same direction, normalized magnitude). Cell (*ii*) reveals a core flaw in the screened Poisson solution: its derivative exceeds our available numerical precision, leading to a vanishing flow field at the center where the solution plateaus. Any cells of this size or larger will exhibit this issue. (**b**) Convergence measured by the average difference at each iteration (maximum normalized to 1) for cells (*i,ii*). Our Eikonal solution converges faster than the other methods by a wide margin at typical cell diameters (*i*).

**Extended Data Fig. 6.**
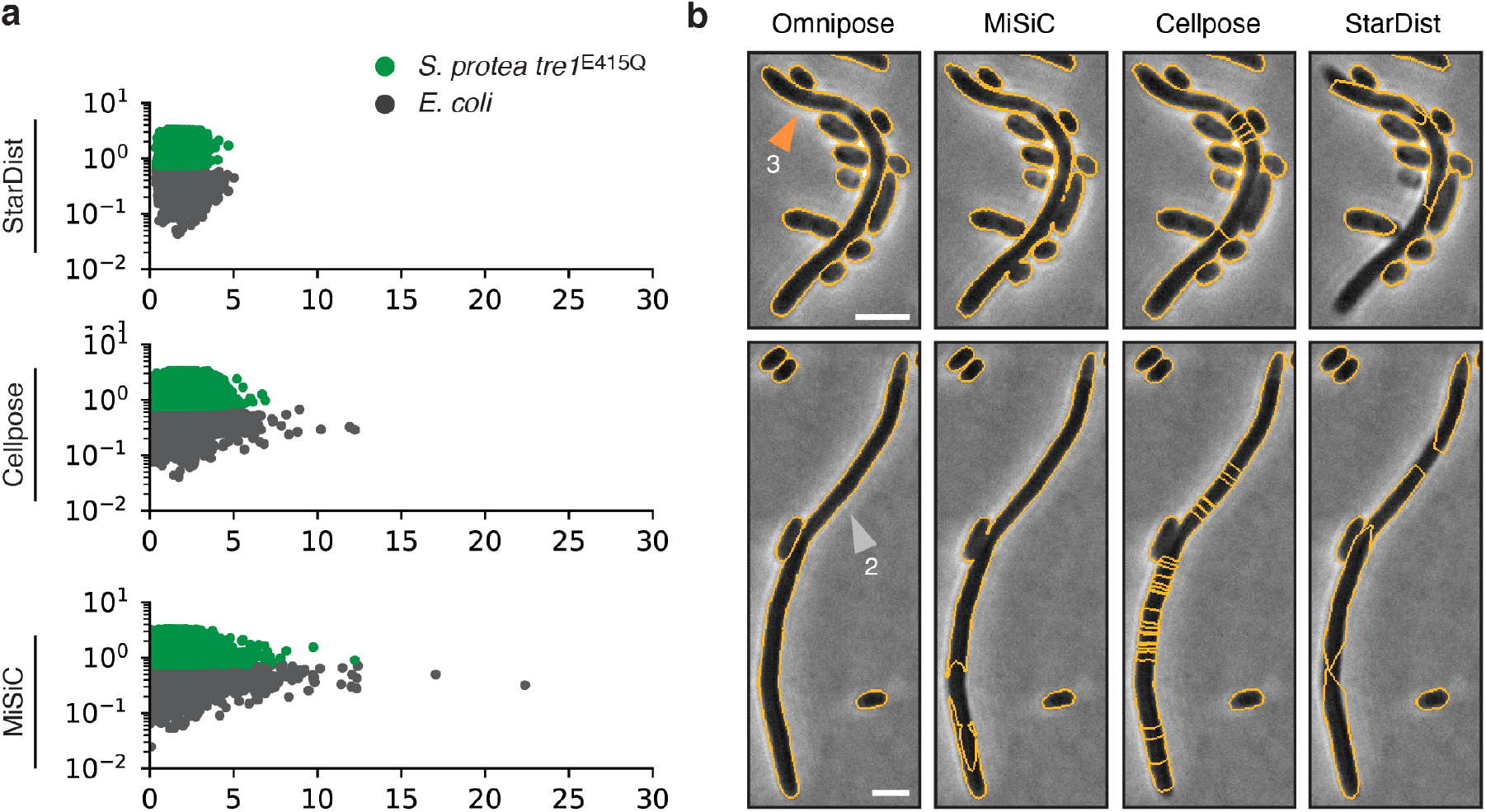
Controls and additional examples. (**a**) Controls segmented by StarDist, Cellpose, and MiSiC. Notably, Cellpose and MiSiC exhibit an enrichment of larger cells even in the control, a consequence of both under-segmented (merged) cells as well as fragments of over-segmented large cells. (**b**) Cells 2 and 3 highlighted in orange and gray plotted in Fig. 5a,d. Scale bars are 1 μm.

**Extended Data Fig. 7.**
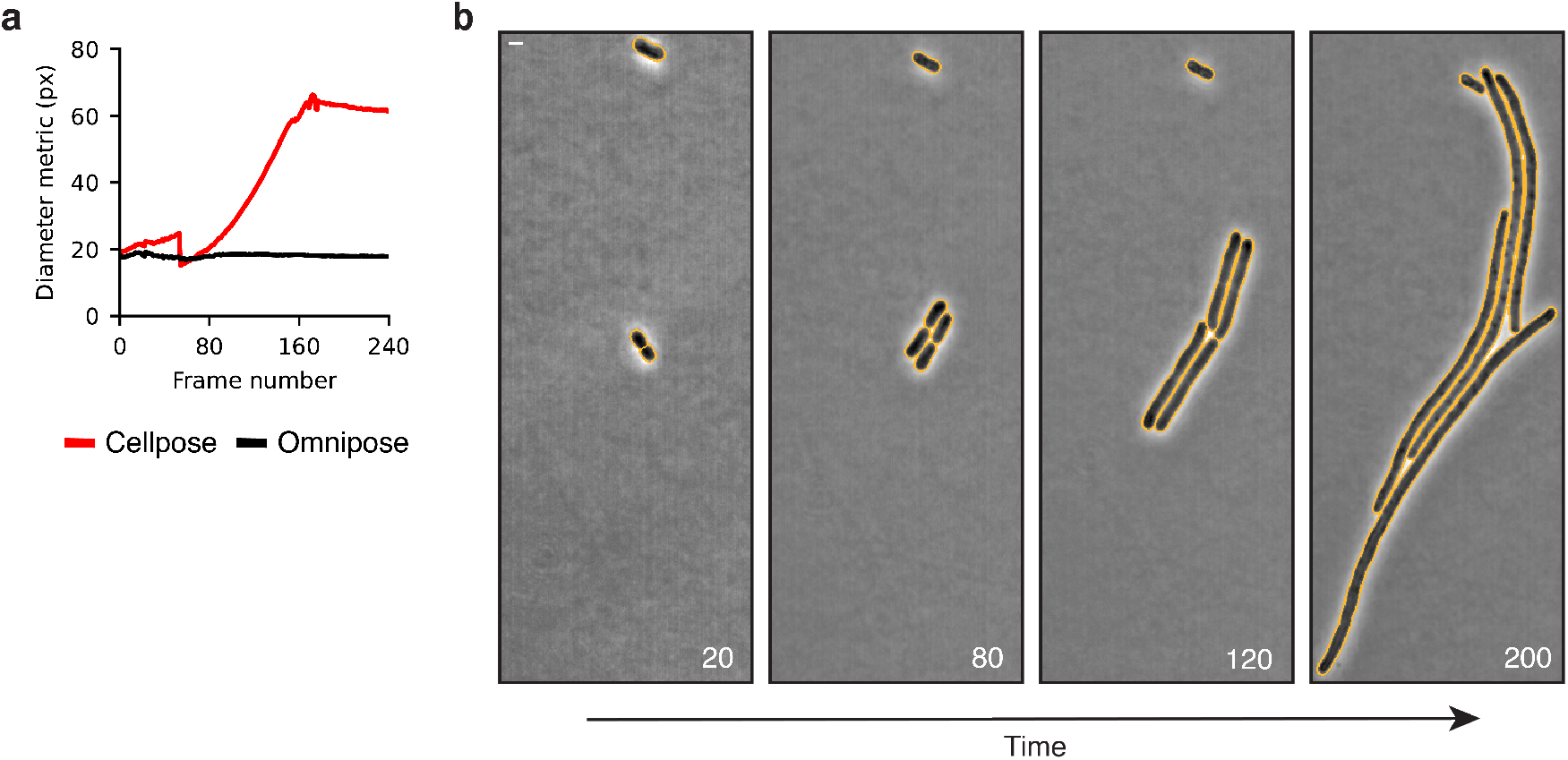
Comparison of diameter metrics on a filamentous microcolony time lapse. (**a**). Cellpose diameter metric is the diameter of the circle with equivalent area. Omnipose diameter metric is proportional to the mean of the distance transform. (**b**) Bacteria displayed are *A. baylyi* transformed with a Δ*ftsN::kan* PCR fragment. Scale bar is 1 μm.

**Extended Data Table 1.**

| Species | Strain | Image count | Cell Count | Cells in GT | Percent of GT | Notes |
| --- | --- | --- | --- | --- | --- | --- |
| *Escherichia coli* | DH5α | 1378 | 98200 | 9733 | 20.6 | Dense microcolonies grown on minimal media. Thin phenotype. ITPG-induced GFP cytosol marker. Time lapse. Imaged by the Wiggins lab. |
|  |  | 141 | 4536 | 4395 | 9.3 | Dense microcolonies on LB. Time lapse. Imaged by the Wiggins lab. |
|  |  | 2 | 2277 | - | - | Treatment with cephalixin. Tn7::GFP. Imaged by the Mougous lab. |
|  | CS703-1<sup>62</sup> | 80 | 23169 | 1299 | 2.6 | Mutant grown on LB and aztreonam. Elongated and branching phenotypes. Time lapse. Imaged by the Mougous lab. |
| *Shigella flexneri* | M90T | 117 | 256618 | 1409 | 3.0 | Treatment with A22. Tn7::GFP. Frames selected from time lapse after 1hr growth. Imaged by the Mougous lab. |
|  |  | 6 | 4482 | 4318 | 9.2 | Treatment with cephalixin. Tn7::GFP. Frames selected from time lapse after 1hr growth. Imaged by the Mougous lab. |
| *Francisella tularensis subsp. novicida* | U112 | 5 | 20166 | 496 | 1.1 | Small and extremely low-contrast cells. Tn7::GFP. Imaged by the Mougous lab. |
| *Acinetobacter baylyi* | ADP1<sup>63</sup> | 2169 | 60601 | 3336 | 7.1 | Deletion of essential gene *murA*. Rounded phenotype. Time lapse. Imaged by the Wiggins lab. |
|  |  | 241 | 1313 | 1133 | 2.4 | Deletion of essential gene *ftsN*. Filamentous phenotype. Time lapse. Imaged by the Wiggins lab. |
|  |  | 540 | 10013 | 2227 | 4.7 | Deletion of essential gene *dnaA*. Filamentous phenotype. Time lapse. Imaged by the Wiggins lab. |
| *Burkholderia thailandensis* | E264<sup>64</sup> | 30 | 62005 | 5122 | 10.9 | Selected panels from a self-intoxication experiment. Cells exhibit internal structure and low contrast in microcolonies. Tn7::GFP. Time lapse. Imaged by the Mougous lab. |
| *Helicobacter pylori* | LHS100<sup>50</sup> | 15 | 13014 | - | - | Helical phenotype. Grown, fixed, and stained with Alexafluor 488 in the lab of |
|  |  |  |  |  |  | Nina Salama. Imaged by the Mougous lab. |
|  |  | 19 | 1668 | 701 | 1.5 | Treated with aztreonam. Filamentous, helical phenotype. Grown, fixed, and stained with Alexafluor 488 in the lab of Nina Salama. Imaged by the Mougous lab. |
| *Caulobacter crescentus* | NA1000<sup>52</sup> | 4 | 1787 | 756 | 1.6 | Grown in HIGG media to induce stalk phenotype. Cultivation and imaging done in the lab of Yves Brun. |
| *Streptomyces pristinaespiralis* | NRRL 2958 | 17 | 2339 | 270 | 0.6 | Grown on rich media to induce filamentous phenotype. Imaged by the Mougous lab. |
| *Vibrio cholerae* | A1552<sup>65</sup> | 2 | 2627 | 2265 | 4.8 | Cells have short but curved morphology and form dense, low-contrast microcolonies. Tn7::GFP. Obtained from the lab of Fitnat Yildiz. Imaged in the Mougous lab. |
| *Serratia proteamaculans*<br>*E. coli* | 568 DH5α | 43 | 100146 | 1244 | 2.6 | 1:1 mixture. *S.p.* labelled via Tn7::GFP, *E.c.* unlabeled. Time lapse. Imaged in the Mougous lab. |
| *Pseudomonas aeruginosa*<br>*Staphylococcus aureus* | PAO1<sup>66</sup><br>USA300 | 3 | 2662 | 3688 | 7.8 | 1:1 mixture. *P.a.* labelled via Tn7::GFP, *S.a.* unlabeled. Imaged in the Mougous lab. |
| *P. aeruginosa*<br>*S. aureus*<br>*V. cholerae*<br>*Bacillus subtilis* | PAO1<br>USA300<br>A1552<br>HM1350 | 21 | 33281 | 4678 | 9.9 | 1:1:1:1 mixture. *P.a.* and *V.c.* labelled via Tn7::GFP, *S.a.* and *B.s.* labelled with red membrane dye. Imaged in the Mougous lab. |
|  |  | 4833 | 700904 | 47070 | 100 |  |

